# Hallmarks of frailty and osteosarcopenia in prematurely aged PolgA^D257A/D257A^ mice

**DOI:** 10.1101/758243

**Authors:** Ariane C. Scheuren, Gommaar D’Hulst, Gisela A. Kuhn, Evi Masschelein, Esther Wehrle, Katrien De Bock, Ralph Müller

**Affiliations:** Institute for Biomechanics, ETH Zurich, Zurich, Switzerland; Laboratory of Exercise and Health, ETH Zurich, Zurich, Switzerland

**Keywords:** aging, osteopenia, sarcopenia, osteosarcopenia, mTORC1, *in vivo* micro-CT

## Abstract

**Background:** Frailty is a geriatric syndrome characterized by increased susceptibility to adverse health outcomes. One major determinant thereof is the gradual weakening of the musculoskeletal system and the associated osteosarcopenia. To improve our understanding of the underlying pathophysiology and, more importantly, to test potential interventions aimed at counteracting frailty suitable animal models are needed.

**Methods:** To evaluate the relevance of prematurely aged PolgA^(D257A/D257A)^ mice as a model for frailty and osteosarcopenia, we quantified the clinical mouse frailty index in PolgA^(D257A/D257A)^ and wild type littermates (PolgA^(+/+)^, WT) with age and concertedly assessed the quantity and quality of bone and muscle tissue. Lastly, the anabolic responsiveness of skeletal muscle, muscle progenitors and bone was assessed.

**Results:** PolgA^(D257A/D257A)^ accumulated health deficits at a higher rate compared to WT, resulting in a higher frailty index at 40 and 46 weeks of age (+166%, +278%, p<0.0001), respectively, with no differences between genotypes at 34 weeks. Concomitantly, PolgA^(D257A/D257A)^ displayed progressive musculoskeletal deterioration such as reduced bone and muscle mass as well as impaired functionality thereof. In addition to lower muscle weights (-14%, p<0.05, -23%, p<0.0001) and fiber area (-20%, p<0.05, -22%, p<0.0001) at 40 and 46 weeks, respectively, PolgA^(D257A/D257A)^ showed impairments in grip-strength and concentric muscle forces (p<0.05). PolgA^(D257A/D257A)^ mutation altered the acute response to various anabolic stimuli in skeletal muscle and muscle progenitors. While PolgA^(D257A/D257A)^ muscles were hypersensitive to eccentric contractions as well as leucine administration, shown by larger downstream signaling response of the mechanistic target of rapamycin complex 1 (mTORC1), myogenic progenitors cultured *in vitro* showed severe anabolic resistance to leucine and robust impairments in cell proliferation. Longitudinal micro-CT analysis of the 6^th^ caudal vertebrae showed that PolgA^(D257A/D257A)^ had lower bone morphometric parameters (e.g. bone volume fraction, trabecular and cortical thickness, p<0.05) as well as reduced remodeling activities (e.g. bone formation and resorption rate, p<0.05) compared to WT. When subjected to 4 weeks of cyclic loading, young but not aged PolgA^(D257A/D257A)^ caudal vertebrae showed load-induced bone adaptation suggesting reduced mechanosensitivity with age.

**Conclusions:** PolgA^(D257A/D257A)^ mutation leads to hallmarks of age-related frailty and osteosarcopenia and provides a powerful model to better understand the relationship between frailty and the aging musculoskeletal system.

## Introduction

Although there is no universally accepted definition of frailty [1], it is considered as an age-related syndrome characterized by the decline of multiple physiological functions, leading to the accumulation of health deficits, and thus a higher vulnerability to adverse health outcomes such as morbidity and mortality [2]. One of the most prominent components of frailty is the progressive weakening of the musculoskeletal system [3–5], leading to common age-related diseases such as osteopenia and sarcopenia. There is growing evidence that both diseases often co-exist in frail older individuals (also termed osteosarcopenia [6, 7]), thereby further increasing the risk for negative outcomes such as falls and fractures [8, 9]. Although several anabolic interventions such as dietary protein supplementation and mechanical stimulation are known to promote muscle and bone formation in young individuals, the molecular insights behind osteopenia and sarcopenia in the elderly population are lacking.

In the field of muscle physiology, studies in humans and rodents have shown that aged muscles are less responsive to well-known anabolic stimuli such as amino acids [10–12] and muscle contractions [13]; this phenomenon, termed “anabolic resistance”, likely results from reduced protein synthesis due to diminished intracellular signaling through the mechanistic target of rapamycin complex 1 (mTORC1) pathway [14–16]. Next to impairments in intra-muscular mTORC1 signaling, age-related sarcopenia has been associated with a decrease in number [17, 18] and proliferation capacity [19, 20] of myogenic progenitors or satellite cells. These are not only instrumental for the maintenance of muscle fibers, but also for the adaptive responses to exercise and regeneration upon injury [21]. In the field of bone physiology, evidence pointing towards altered mechanosensitivity with age has also been shown in humans [22] and in mice [23–26]. However, this effect might be site-specific as studies using a tibia-loading model showed a reduced response of trabecular [23, 24] and cortical [25, 26] bone formation with age, while bone adaptation in response to loading of the caudal vertebrae was maintained with age [27].

Therefore, whether and how age-related changes in the responsiveness to anabolic stimuli occur remains unclear. A better understanding of the pathophysiology of osteosarcopenia will help to identify interventions to strengthen the musculoskeletal system, which ultimately will be beneficial for the prevention and/or treatment of frailty.

In order to address this, tools such as the frailty index (FI) have been established to quantify the accumulation of age-related health deficits (e.g., loss of hearing, tremoring, comorbid diseases) in humans [28] and more recently, also in mice [29]. Indeed, the striking similarities between key features of the FI scores in humans and in mice [30] have highlighted the potential of rodent frailty models to not only improve our understanding of frailty but also serve as a tool to test responses to interventions designed to modify (or even prevent) frailty [31–33]. In this study, we aimed to evaluate the PolgA^(D257A/D257A)^ mouse (referred to as PolgA), which due to elevated mitochondrial DNA point mutations and systemic mitochondrial dysfunction, exhibits an accelerated aging phenotype [34, 35], as a model of frailty and osteosarcopenia. While these mice are known to develop multiple signs of aging (e.g., hair loss, greying, hearing loss) earlier (around 40 weeks of age) than their wild type littermates (PolgA^+/+^, referred to as WT), the frailty phenotype has to the best of our knowledge not yet been assessed in these mice. Furthermore, although several studies have reported lower muscle weights in PolgA mice compared to their WT littermates [34, 36, 37], little is known about their muscle quality and functionality. To address this, forelimb grip-strength and concentric muscle forces were measured *in vivo* and *ex vivo*, respectively, in addition to the evaluation of hind limb muscle masses. Furthermore, the response to acute anabolic stimuli such as eccentric contractions and the leucine administration were assessed. With respect to the bone phenotype, only two studies have reported reduced femoral bone density using X-ray densitometry [34, 35]. Although this technique is still the gold-standard to assess bone mineral density (i.e. bone quantity) clinically in humans, it does not provide insight regarding the quality of bone tissue. Therefore, bone phenotyping in pre-clinical studies is commonly performed using high-resolution micro-computed tomography (micro-CT) as it allows additional standardized evaluation of the three-dimensional bone microarchitecture [38]. Furthermore, using *in vivo* micro-CT, dynamic bone remodeling activities can be tracked longitudinally providing information both on bone formation and bone resorption [25, 39]. These markers are important for better understanding of age-related changes in bone microarchitecture due to osteopenia. Coupled with longitudinal measurements of FI, we aimed to monitor individual mice during the process of aging, allowing us to capture the onset of osteosarcopenia and to track other signs of aging. Lastly, by longitudinally monitoring bone adaptation in response to a long-term mechanical loading intervention, we aimed to investigate whether PolgA bones show altered mechanosensitivity with age.

## Materials and Methods

### Study Design

The study consisted of three parts. For parts 1 and 2, female mice were aged up to 46 weeks and divided into four groups. Three of the groups were used to cross-sectionally compare the musculoskeletal and frailty phenotype of PolgA and WT mice at 34, 40 and 46 weeks, respectively (*in vivo* and *ex vivo*, part 1). The fourth group was longitudinally monitored between the ages of 20 and 40 weeks to investigate the changes in bone microarchitecture and frailty over time (*in vivo*, part 2). Lastly, the effects of various anabolic interventions on bone and muscle tissue were assessed (*in vivo*, *ex vivo*, *in vitro*). The number of mice and sample sizes in various animal studies are provided directly in the figure legends and in table S1. The sample sizes for *in vivo* experiments were selected based on previous experiments and power analysis (power set at 0.80) was used to determine the sample size required for *ex vivo* experiments. All mouse experiments described in the present study were carried out in strict accordance with the recommendations and regulations in the Animal Welfare Ordinance (TSchV 455.1) of the Swiss Federal Food Safety and Veterinary Office and results are reported following the principles of the ARRIVE guidelines (www.nc3rs.org.uk/arrive-guidelines).

### Animals

All animal procedures were approved by the local authorities (licence numbers 262/2016 and ZH255-16, Verterinäramt des Kantons Zürich, Zurich, Switzerland). A colony of the mouse strain expressing an exonuclease-deficient version of the mitochondrial DNA polymerase γ (PolgA^D257A^, B6.129S7(Cg)-Polg^tm1Prol^/J, JAX stock 017341, The Jackson Laboratory) was bred and maintained at the ETH Phenomics Center (12h:12h light-dark cycle, maintenance feed and water ad libitum, 3-5 animals/cage). The mice were bred and genotyped as described in Supplementary Materials (SM). In order to confirm that the premature aging phenotypes were associated with mitochondrial dysfunction, the activity of complex IV enzyme (COX IV) in *m. gastrocnemius* (GAS) was measured (SM). Compared to WT, PolgA muscles had lower COX IV activity both at 40 and 46 weeks (-19% and -24%, p<0.0001, respectively), thus confirming that the mice used in this study had the same phenotype as those previously reported [36, 37, 40].

### Quantification of Frailty Index (FI)

As recommended in the recently established toolbox for the longitudinal assessment of healthspan in aging mice [41], frailty was quantified using the Mouse Clinical FI [29], which includes the assessment of 31 non-invasive clinical items. For 29 of these items, mice were given a score of 0 if not present, 0.5 if there was a mild deficit, and 1 for a severe deficit. The final two items were weight and body surface temperature, which were scored based on the number of standard deviations from a reference mean in young adult mice as previously described [29]. To compare rates of deficit accumulation between PolgA and WT, the natural log of the FI was plotted against age. The slope of this line provides an estimate of the rate of deficit accumulation, as shown in previous studies [29, 42, 43].

### Forelimb Grip-Strength

Forelimb grip-strength was measured using a force tension apparatus (Grip Strength Meter, 47200, Ugo Basile) at 40 and 46 weeks of age. Once mice gripped the stationary bar with their forepaws, they were pulled horizontally at the base of their tail until they let go of the bar. The process was repeated 5 times to determine the average peak grip force value (gram-force) used for analysis. All measurements were performed by the same experienced user.

### Muscle harvesting and force measurements

PolgA and WT GAS, TA and *m. soleus* (SOL) were excised under anesthesia (5% isoflurane/oxygen) and snap frozen in liquid nitrogen. Furthermore, EDL from both legs were dissected and maintained in 4°C Krebs–Henseleit buffer supplemented with 1×MEM amino acid mixture (Invitrogen) and 25mM glucose. For assessment of *ex vivo* force production, the EDL of the right leg was attached to the lever arms of an Aurora system (Aurora Scientific) and submerged in continuously gassed Krebs–Henseleit buffer maintained at 37°C. After 5 min of temperature acclimation, muscle length was adjusted until a single stimulus pulse elicited maximum force during a twitch (Lo) under isometric conditions. After 5 min rest, a force frequency protocol was initiated by subsequently providing a pulse train (lasting 250 ms) of 1-30-50-80-150-250 and 300Hz with 1 min rest between every intensity.

### Eccentric contractions (ECC) protocol

5 min after the force frequency protocol, EDL from the right legs were subjected to an eccentric training protocol according to O’Neil et al. [44] consisting of 60 contractions in a 22 min time window. In this study, the pulse train was changed to 200Hz (compared to 100Hz in O’Neil et al.), because pilot studies suggested that 100Hz was not sufficient to induce maximal eccentric force in older muscle. After the contractions, the muscle was maintained in 37°C Krebs–Henseleit buffer + 1xMEM amino acid mixture and 25mM glucose before snap-freezing 1h after the last contraction. The control EDL from the contralateral leg was kept in the same 37°C Krebs–Henseleit buffer during the complete ECC period.

### Leucine administration

46-week-old mice were fasted for 5h in the beginning of their light cycle, after which saline (0.9% NaCl, CTL) or leucine (0.4g/kg, LEU) was administered via oral gavage. L-Leucine (Sigma Aldrich) was dissolved in a stock solution of 40g/L, heated (40-50°C) and acidified as described previously [45]. 30 min later, hind limb muscles were weighted and snap frozen for further analysis.

### Leucine stimulation in vitro

Primary muscle progenitor cells (MPs) were isolated from muscle tissue as described in SM. 200’000 cells were seeded in 6-well-plates. Myogenic differentiation medium containing low-glucose DMEM, 2% Horse-serum (HS) (Invitrogen) and 1% P/S was added 24h later. 72h later, fully differentiated MPs were treated according to one of the following conditions: (1) STV; 1h DMEM w/o amino acids (Biomol GmbH) + 10% dialyzed FBS (Invitrogen) + 1% P/S, (2) LEU 0.8; 1h STV + 1h 0.8mM L-Leucine (Sigma Aldrich) and (3) LEU 5; 1h STV + 1h 5mM L-Leucine. Experiments were performed in biological triplicate and technical duplicate. Edu analysis was performed according to manufacturer’s protocol (Thermo Fischer Scientific) after a 4h pulse with 1µg/ml Edu. For proliferation analysis, 20’000 cells were plated on 12-well-plates, and 3 plates were counted per time point over 5 days.

### Immunoblotting

Details have been described previously [46]. Briefly, bone tissue (20-25mg) was first pulverized with mortar and pestle and subsequently homogenized with a tissue homogenizer (Omni THq). Muscle tissue (10-15 mg) was homogenized directly with a tissue homogenizer. Lysis was performed in ice-cold lysis buffer (for specifications see SM, Table S2). Homogenates were centrifuged at 10’000 g for 10 min at 4°C. Supernatant was collected and protein concentration was measured using the DC protein assay kit. 10-25 µg of total protein was loaded in a 15-well pre-casted gradient gel (Bio-rad, 456-8086). After electrophoresis, a picture of the gel was taken under UV-light to determine protein loading using strain-free technology. Proteins were transferred via semi-dry transfer onto a PVDF membrane (Bio-rad, 170-4156) and subsequently blocked for 1 h at room temperature with 5% milk in TBS-Tween. Membranes were incubated overnight at 4 °C with primary antibodies (listed in SM, Table S2). The appropriate secondary antibodies for anti-rabbit and anti-mouse IgG HRP-linked antibodies (SM, Table S2) were used for chemiluminescent detection of proteins. Membranes were scanned with a chemidoc imaging system (Bio-rad) and quantified using Image lab software (Bio-rad).

### Micro-CT imaging and analysis

For analysis of femoral, the right femora were harvested, placed in 70% Ethanol and scanned with micro-CT (micro-CT40, Scanco Medical AG, isotropic nominal resolution: 10 µm; 55 kVp, 145 µA, 200 ms integration time). A 3D constrained Gaussian filter (sigma 1.2, support 1) was applied to the image data, after which images were segmented by a global thresholding procedure [47]. Standard bone microstructural parameters were calculated in trabecular (BV/TV, Tb.Th, Tb.N, Tb.Sp), cortical (Ct.Ar/Tt.Ar, Ct.BV, Ct.MV, Ct.Ar, Tt.Ar, Ct.Th, Ps.Pm, Ec.Pm) and whole (AVD, length) bone using automatically selected masks for these regions [48].

For analysis of the 6^th^ caudal vertebra (CV6), *in vivo* micro-CT (vivaCT 40, Scanco Medical AG, isotropic nominal resolution: 10.5 µm; 55 kVp, 145 µA, 350 ms integration time, 500 projections per 180°) was performed every 2 weeks between 20 and 40 weeks of age. Animals were anesthetized with isoflurane (induction/maintenance: 5%/1-2% isoflurane/oxygen). Micro-CT data was processed and standard bone microstructural parameters were calculated in trabecular, cortical and whole bone by using automatically selected masks for these regions as described previously [49].

To calculate dynamic morphometric parameters, micro-CT images from consecutive time-points were registered onto one another. The voxels present only at the initial time point were considered resorbed whereas voxels present only at the later time point were considered formed. Voxels that were present at both time points were considered as quiescent bone. By overlaying the images, morphometrical analysis of bone formation and resorption sites within the trabecular region allowed calculations of bone formation rate (BFR), bone resorption rate (BRR), mineral apposition rate (MAR), mineral resorption rate (MRR), mineralizing surface (MS) and eroded surface (ES) [39].

### Analysis of bone turnover markers

Immediately following euthanasia, blood samples from 40-week-old PolgA and WT mice were obtained by cardiac puncture. Serum was separated by centrifugation and stored at -80°C until further analysis. Markers for bone formation and resorption were measured in technical duplicates using ELISA kits for N-terminal propeptide of type I procollagen (PINP, AC-33F, Immunodiagnostics) and for C-terminal cross-linked telopeptides of type I collagen (CTX-I, RatLaps, AC-06F1, Immunodiagnostics) according to manufacturer’s instructions.

### Cyclic mechanical loading of CV6

CV6 were subjected to a cyclic loading regime, which has previously been shown to have anabolic effects in 15-week-old female C57BL/6J mice [50]. Briefly, stainless steel pins (Fine Science Tools) were inserted into the 5^th^ and 7^th^ caudal vertebrae of 35- and 12-week-old female mice. Three weeks after surgery, the mice received either sham (0N control) or 8N cyclic (10 Hz) loading for 5 minutes, 3x/week over 4 weeks. Weekly *in vivo* micro-CT images (vivaCT 40 or vivaCT80, Scanco Medical AG) were acquired and analyzed as described above.

### Statistical analysis

Data is reported as mean and standard error of the mean (±SEM), unless otherwise stated. All analysis, with the exception of the longitudinal micro-CT data, was performed using GraphPadPrism (8.0.0). Unpaired or paired Student’s t-test, Mann Whitney test, one- or two-way ANOVA with post hoc multiple comparison testing were performed as indicated in figure legends. For the analysis of the longitudinal micro-CT data, linear mixed-effects modelling was used with Tukey’s post hoc multiple comparison testing corrected with Bonferroni criteria (SPSS 24.0.0.0). Fixed-effects were allocated to the age and genotype. Random-effects were allocated to the animal to account for the natural differences in bone morphometry in different mice. Significance was set at α<0.05 in all experiments.

## Supplementary Materials

Materials and Methods

Table S1. Animal experiments: Overview, setup and analysis details.

Table S2: Reagents and antibodies used for immunoblotting.

Table S3: Cortical bone morphometry of femora.

## Results

### With age, PolgA mice become frailer and display signs of co-existing osteopenia and sarcopenia

To characterize the frailty and the musculoskeletal phenotypes of the PolgA model, female PolgA^(D257A/D257A)^ (referred to as PolgA) and wild type littermates (PolgA^(+/+)^, referred to as WT) were aged in parallel and sacrificed at 34, 40 and 46 weeks, respectively. Using the clinical mouse frailty index (FI) as a tool to quantify the accumulation of health deficits [29], we observed that older (40 and 46 weeks), but not younger (34 weeks) PolgA mice had higher FI scores compared to WT (*Figure* 1A). This divergence between genotypes at later time-points was also observed when FI was assessed longitudinally in individual mice at 34, 38 and 40 weeks, respectively. Specifically, between 34 and 40 weeks, the mean FI in PolgA mice increased from 0.05±0.02 to 0.15±0.02 (mean±SD, p<0.0001), whereas the increase in WT from 0.04±0.03 to 0.06±0.04 was less pronounced (p<0.05, *Figure* 1B). Thus, compared to WT, PolgA mice had 73% (p<0.05) and 128% (p<0.001) higher FI at 38 and 40 weeks, respectively.

**Fig. 1.**
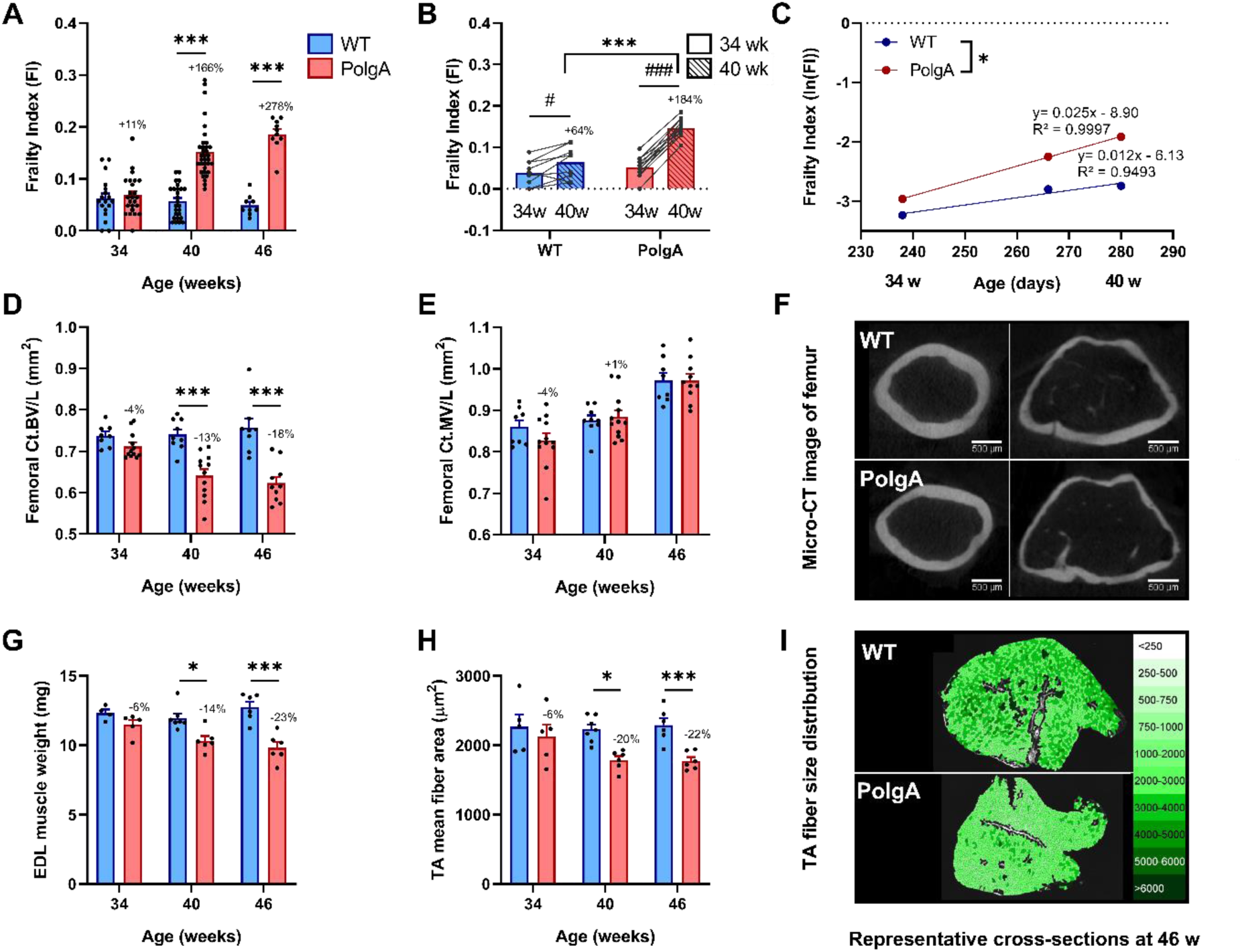
Comparison of the frailty and musculoskeletal phenotypes at different ages. (**A**) Frailty Index (FI) assessed in PolgA and WT at 34, 40 and 46 weeks (n=9-35/group). (**B**) Longitudinal monitoring of FI in individual mice (n=9-12/group) at 34 (solid bars) and 40 weeks (striped bars). (**C**) The natural logarithm FI was plotted as a function of age. The slope of the regression lines through these data, which represent the rate of deficit accumulation, was higher in PolgA mice. (**D-I**) PolgA mice displayed lower bone and muscle mass and cross-sectional area compared to WT. (**D**) Femoral cortical bone volume (Ct.BV) and (E) cortical marrow volume (Ct.MV) (n=8-12/group). (G) Muscle weight and (H) fiber area (n=5-7/group). Representative cross-sections of (C) femoral bone and (F) m. tibialis anterior (TA) muscle of PolgA and WT mice at 46 weeks. (Data represent mean±SEM; * p<0.05; **p<0.01, ***p<0.0001 WT (blue bars) vs PolgA (red bars) determined by unpaired t-test, corrected for multiple comparisons by Bonferroni; for (**B**), ***p<0.0001 WT vs PolgA and #p<0.05, ###p<0.0001 over time determined by two-way ANOVA)

Furthermore, the slope of the natural logarithm of the FI versus age curve, which has been shown to correspond to the rate of deficit accumulation in humans [43], was 0.025 in PolgA mice and 0.012 in WT, suggesting that PolgA mice became frailer faster than WT (p<0.05, *Figure* 1C).

Concomitant to the increase in FI with age, older PolgA mice displayed deteriorations of the musculoskeletal system. Specifically, micro-CT analysis of the femora showed that the cortical bone volume (Ct.BV) was similar between genotypes at 34 weeks of age (p>0.05), but then diverged over time such that PolgA had lower Ct.BV at 40 and 46 weeks, respectively (p<0.0001, *Figure* 1D). In line with the decline in Ct.BV, the cortical area (Ct.Ar) and thickness (Ct.Th) were lower in PolgA femora compared to WT at 40 (-13% and -11%, p<0.0001) and 46 weeks (-17% and -15%, p<0.0001), respectively (Table S3). The cortical marrow volume (Ct.MV) increased both in WT and PolgA with age with no differences between genotypes at any of the ages (p>0.05, *Figure* 1E). The total cross- sectional area within the periosteal envelope (Tt.Ar) was similar between genotypes at 34 weeks of age, but significantly lower in PolgA compared to WT at 40 and 46 weeks (-5% and -7%, p<0.01), respectively (Table S3). Hence, while PolgA and WT showed similar endocortical expansion with age, PolgA showed reduced periosteal expansion at 40 and 46 weeks, respectively. Furthermore, PolgA had slightly but significantly shorter femora compared to WT at 40 and 46 weeks (-2% and -1%, p<0.001, Table S3), respectively. Regarding the muscle tissue, the weights of *m. extensor digitorum longus* (EDL) and fiber cross-sectional area of *m. tibialis anterior* (TA) were similar between genotypes at 34 weeks (p>0.05), but lower in PolgA mice at 40 (-14% and -20%, p<0.01) and 46 weeks (-23% p<0.0001, -22% p<0.01), respectively (p<0.05, *Figure* 1G-I). Similarly, lower weights of TA and *m. gastrocnemius* (GAS) in PolgA mice as compared to WT were observed at 40 (-20%, p<0.0001 and -23%, p<0.0001) and 46 weeks (-22%, p<0.0001 and -16%, p<0.05), respectively.

### Decreased forelimb grip-strength and concentric force in PolgA mice

Considering that PolgA mice showed clear reductions in muscle mass and cross-sectional area at 40 and 46 weeks, we aimed to comprehensively characterize the muscular functionality at these time-points. At 40 weeks, *in vivo* assessment of the forelimb grip-strength, a widely used method to assess muscle function in rodents, showed that PolgA mice had lower grip-strength relative to bodyweight (p<0.05, *Figure* 2A). Furthermore, when the EDL was subjected to a force-frequency protocol, PolgA showed a tendency towards decreased absolute force at 250 and 300Hz (p<0.10), while relative force was not affected (*Figure* 2B,C). At 46 weeks, the grip-strength was 11% lower in PolgA mice compared to WT, but did not reach significance (*Figure* 2D). The differences in forces however, were more exacerbated with a lower absolute and relative force at 80 to 300Hz (absolute) and 250 to 300Hz (relative) in the PolgA (p<0.05, *Figure* 2E,F).

**Fig. 2.**
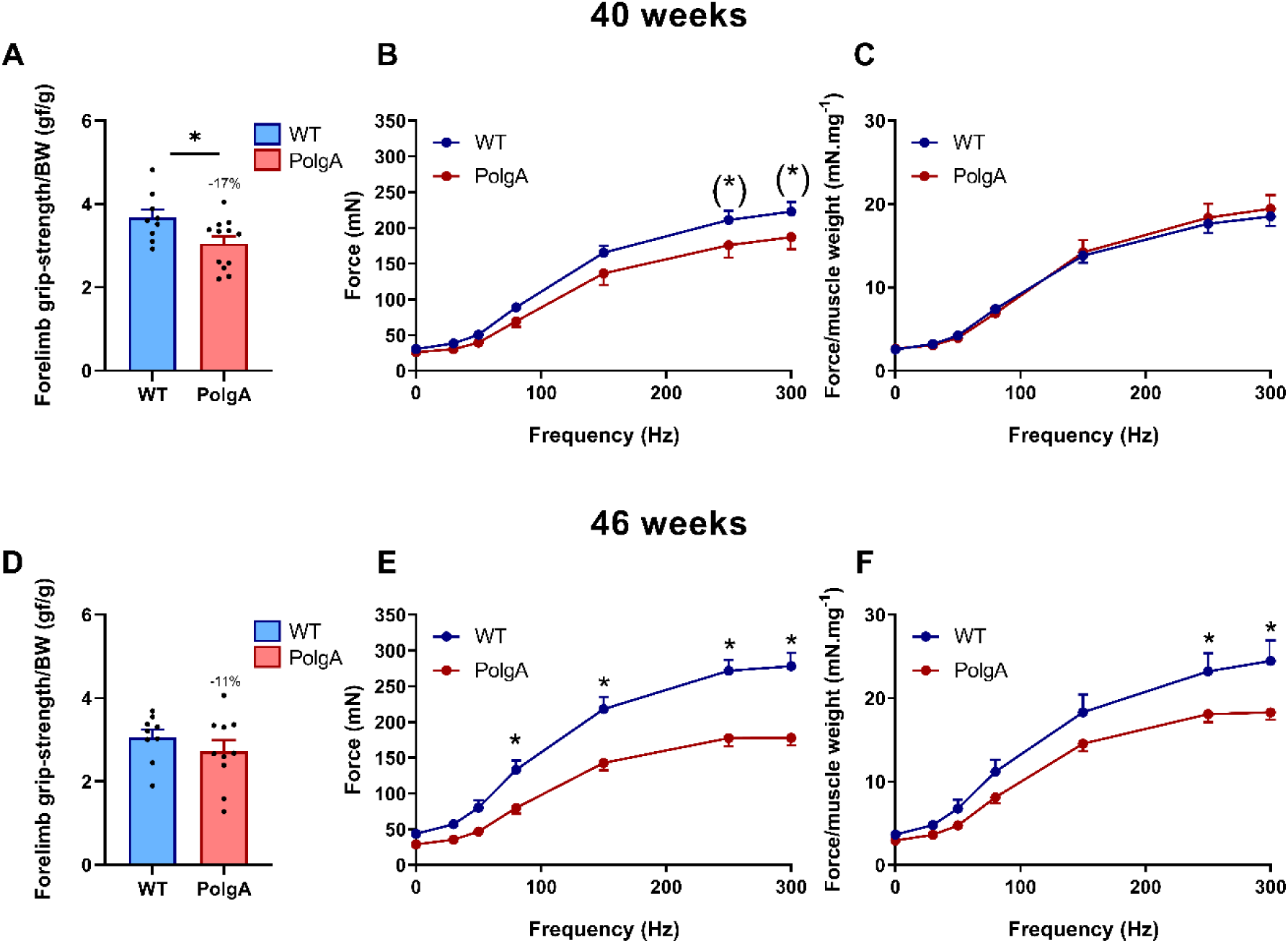
Muscle phenotyping at 40 (upper row, **A**-**C**) and 46 weeks (lower row, **D-F**). (**A**,**D**) Forelimb Grip-Strength (n=9-12/group). (**B**,**E**) Force Frequency at 1, 30, 50, 80, 150, 250, 300 Hz. (**C**,**F**) Force frequency relative to EDL muscle weight (n=6-8/group). (Data represent mean±SEM; (*)p<0.10, *p<0.05 by student t-tests (**A**-**D**) and two-way ANOVA (**E**-**H**))

### PolgA mutation induces higher basal and eccentric contraction (ECC) evoked mTORC1 and mechanotransduction signaling

Because PolgA muscles were atrophied and weaker than those of their WT littermates of the same chronological age, we used an isolated *ex vivo* model to assess the acute response to ECC, which have been shown to effectively activate mTORC1, the main regulator of skeletal muscle protein synthesis [51, 52]. Data of muscle forces evoked by ECC are presented in table 1. PolgA had similar average force during the 60 ECC contractions, but tended to have a lower peak force during the first set (table 1).

**Table 1.** Forces eccentric muscle contractions at 40 and 46 weeks. (^(^*^)^p<0.10, **p<0.001, ***p<0.0001 WT vs PolgA at 40 and 46 weeks, respectively determined by unpaired t-test, corrected for multiple comparisons by Bonferroni)

|  | **40 weeks** |  | **46 weeks** |  |
| --- | --- | --- | --- | --- |
| Average peak force (mV) | WT | PolgA | WT | PolgA |
| all contractions | 149.9±9.7 | 142.8±15.7 | 141.6±9.6 | 137.3±8.8 |
| 1<sup>st</sup> set | 193.3±12.4 | 158.8±17.5 | 188.5±6.6 | 164.6±8.6(*) |
| 10<sup>th</sup> set | 108.8±8.0 | 107.5±11.2 | 98.6±8.2 | 111.8±5.8 |

At 40 weeks, ECC evoked increased phosphorylation of the downstream mTORC1 kinase Ribosomal protein S6 Kinase 1(S6K1) at Thr389 compared to the contralateral control leg in both WT (+147±50%, p<0.05) and PolgA (406±161%, p<0.05), but the increase was more pronounced in PolgA resulting in a higher pS6K1 upon ECC in the PolgA (Δ_ECC-CTL_, p<0.05) (*Figure* 3A). The direct downstream kinase of pS6K1, S6 Ribosomal Protein (RPS6) showed similar increases in phosphorylation at Ser235/236 upon ECC in WT and PolgA (64±24% and 352±146%, p<0.05), whereas there was only a trend towards higher activation in PolgA compared to WT (Δ_ECC-CTL_, p=0.07) (*Figure* 3B). At 46 weeks, downstream mTORC1 targets showed similar effects with a trend towards hyper-phosphorylated pS6K1 and pRPS6 after ECC contractions in the PolgA compared to WT (p=0.10, *Figure* 3D,E). Moreover, basal pRPS6 was higher in PolgA (1257±427%, p<0.05) compared to WT (*Figure* 3E), suggesting both basal and contraction-induced hyper activation of mTORC1 in PolgA mutated muscle.

**Fig. 3.**
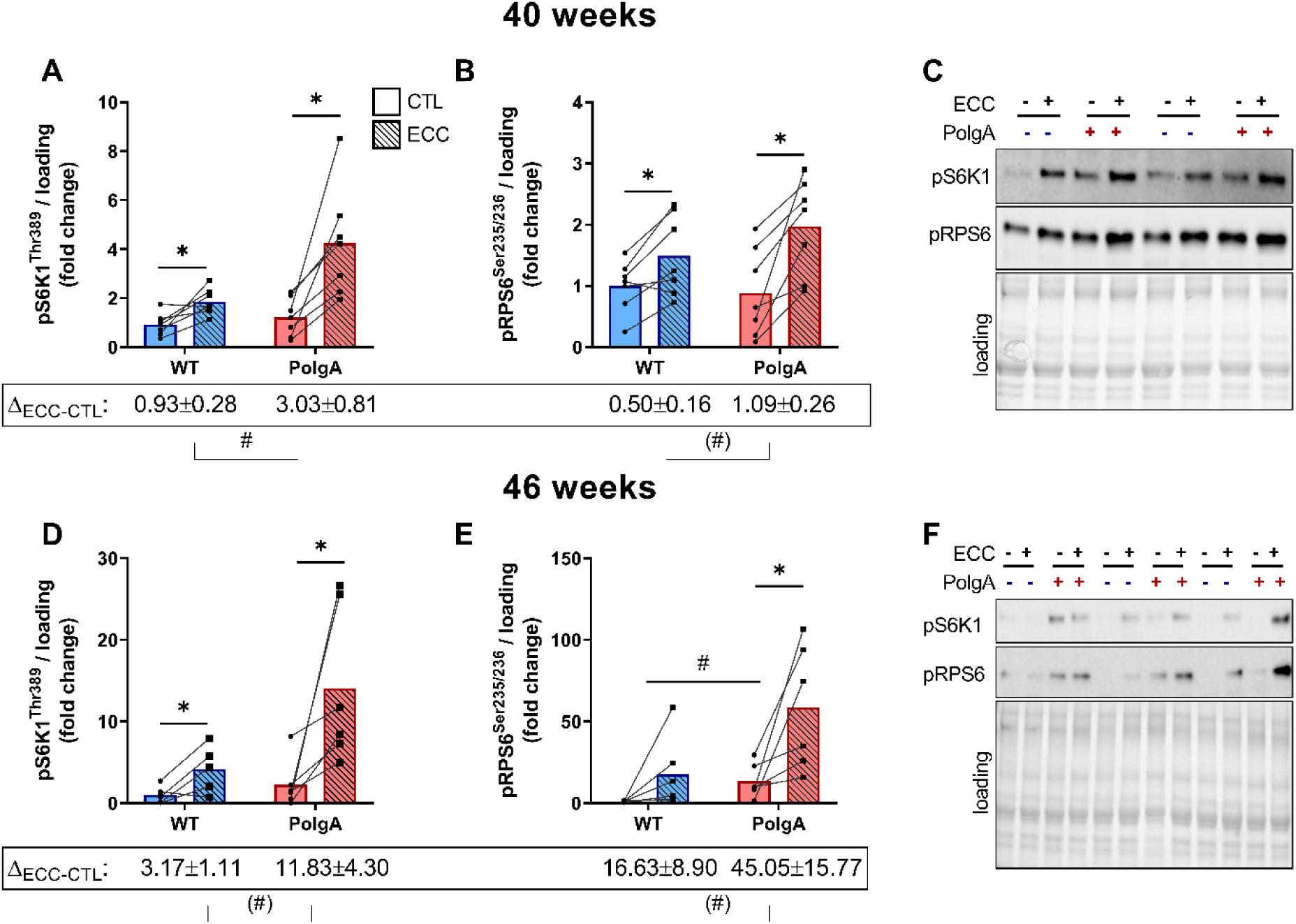
Downstream mTORC1 signaling upon eccentric contraction (ECC) in WT and PolgA EDL. Downstream mTORC1 targets pS6K1 and pRPS6 increased upon ECC both at 40 wk (**A**,**B**) and 46 wk (**D**,**E**). (**C**,**F**) representative blots. (Data represent mean±SEM, n=6-7/group, within genotypes: *p<0.05, **p<0.001 ECC (squares/striped bars) vs. CTL (circles/solid bars) by paired student’s t-test, between genotypes: (#)p<0.10, #p<0.05 by unpaired students t-test ΔECC-CTL and basal values)

One of the proposed pathways for high-load contractions to regulate mTORC1 is via activation of the stress responsive mitogen-activated protein kinase (MAPK) pathway [53]. To investigate whether the hyperactive mTORC1 signaling in the PolgA muscle was related to increased MAPK response, we measured activation of (Stress-Activated Protein Kinase/Jun-amino-terminal Kinase) pSAPK/JNK upon ECC. At 40 weeks, the stress-responsive pSAPK/JNK at Thr138/Tyr185 after ECC increased by 113±35.8% (p<0.05) in WT and by 156±26% (p<0.001) in PolgA (*Figure* 4A), with a tendency towards hyperactivation in PolgA compared to WT (Δ_ECC-CTL_, p=0.09). At 46 weeks, the increase in pSAPK/JNK upon ECC was more pronounced in the PolgA (2410±1035%, p<0.05) than in the WT (1410±179%, p<0.05) resulting in a higher phosphorylation in PolgA muscle after ECC (Δ_ECC-CTL_, p<0.05) and a tendency towards higher basal pSAPK/JNK (p=0.07, *Figure* 4D,J). Interestingly, another arm of the stress-responsive MAPK pathway, p42-44 MAPK at Thr202/Tyr204, was not affected by ECC; however, basal p44/42 signaling was higher in PolgA muscle compared to WT at 46 weeks (*Figure* 4B,D). These data show that hyperactive mTORC1 signaling mirrors increased basal and contraction-induced MAPK signaling in PolgA mutated muscle.

**Fig. 4.**
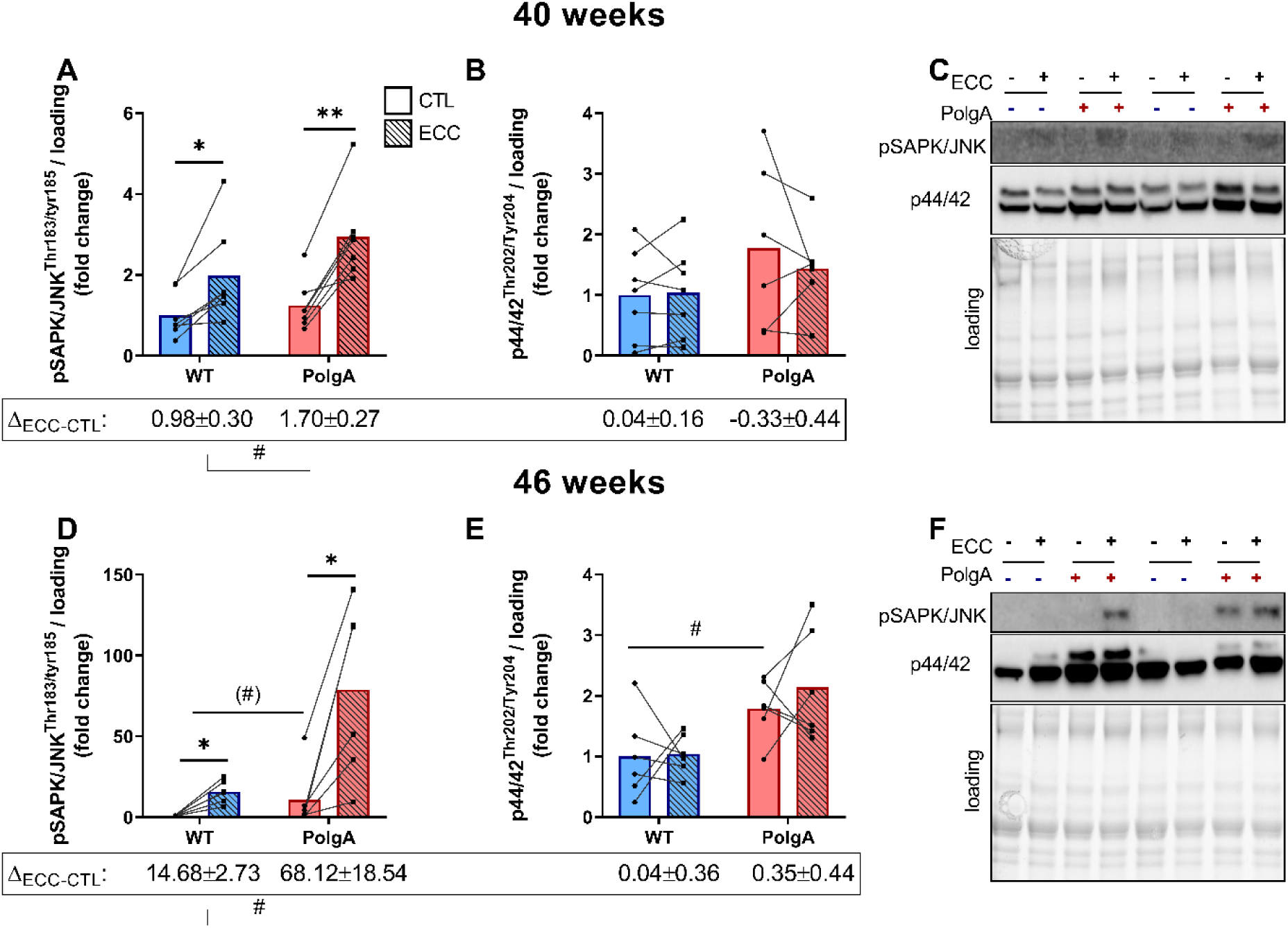
Downstream MAPK signaling upon eccentric contraction (ECC) in WT and PolgA EDL. pSAPK/JNK, but not p44/42 MAPK ERK1/2 increased upon ECC both at 40 wk (**A**,**B**) and 46 wk (**D**,**E**). (**C**,**F**) representative blots. (Data represent mean±SEM, n=6-7/group, within genotypes: *p<0.05, **p<0.001 ECC (squares/striped bars) vs. CTL (circles/solid bars) by paired student’s t-test, between genotypes: (#)p<0.10, #p<0.05 by unpaired students t-test ΔECC-CTL and basal values)

### PolgA whole muscles are hypersensitive to leucine in vivo, while their primary myotubes are resistant to leucine in vitro

To examine whether other anabolic stimuli also hyper activate mTORC1 signaling in PolgA muscle, we administered a submaximal (0.4g/kg) dose of leucine via gavage to 46-week-old WT and PolgA littermates. Leucine induced an increase in pRPS6 in both WT (p<0.05) and PolgA (p<0.0001) TA, whereas pS6K1 was also increased in PolgA (p<0.001). The increase was 3.3±0.8 and 11.4±5.3 fold higher in PolgA for both pRPS6 and pS6K1, respectively (p<0.001, *Figure* 5A-C). These data demonstrate that the amino acid leucine increases downstream mTORC1 signaling to a greater extent in PolgA mutated compared to WT muscle.

**Fig. 5.**
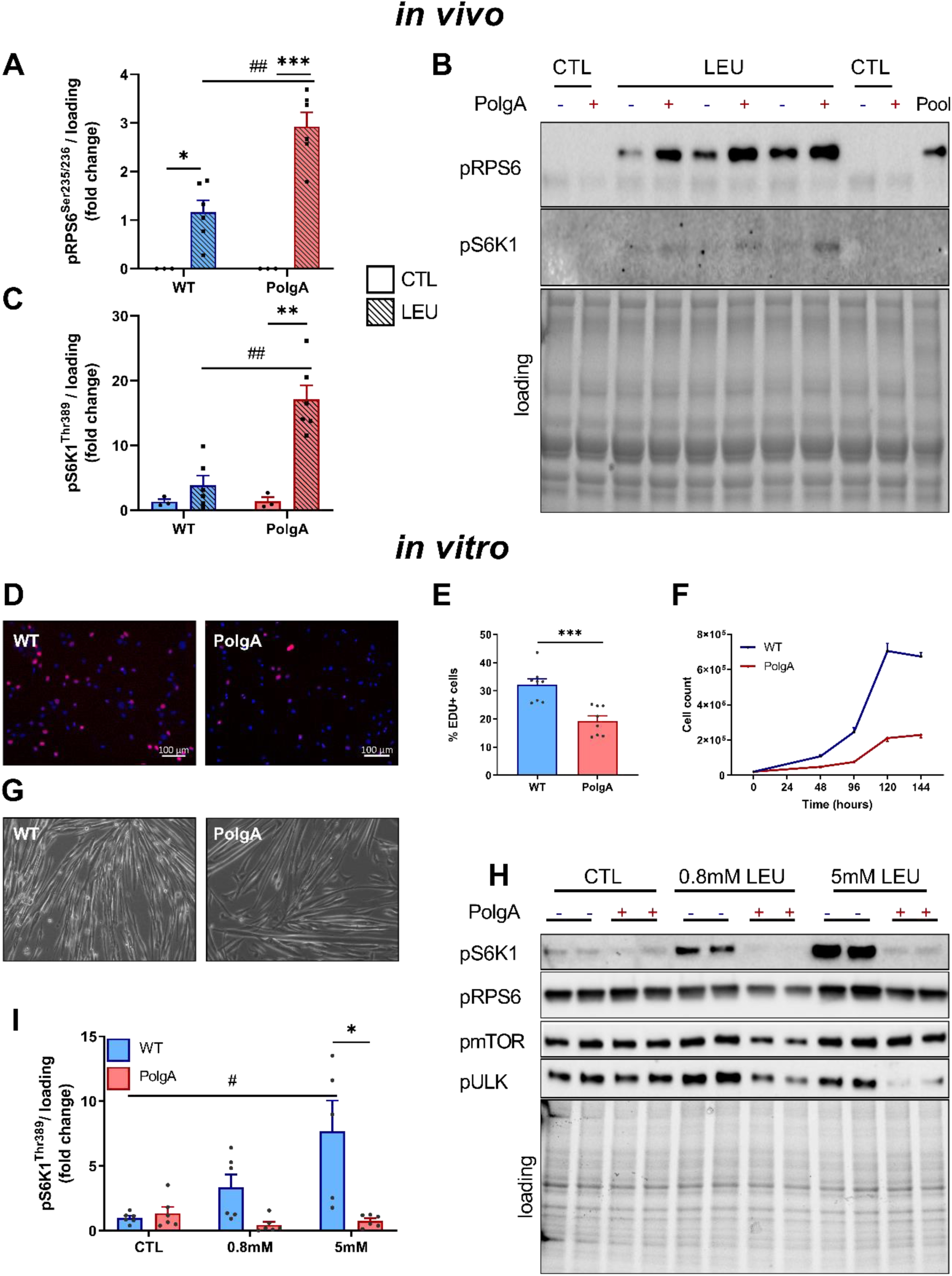
Downstream mTORC1 signaling after leucine administration *in vivo* and *in vitro*. (**A**-**C**). pRPS6 and pS6K1 signaling 30 min after submaximal dose of leucine gavage *in vivo* in 46-week-old WT and PolgA (Data represent mean±SEM, n=3-6/group, within genotypes: *p<0.05, **p<0.001, ***p<0.0001 LEU (squares/striped bars) vs. CTL (circles/solid bars), between genotypes: (#)p<0.10, #p<0.05, ##p<0.001 determined by two-way ANOVA). (**C**) Representative blots. (**D**-**I**) Proliferation and downstream mTORC1 signaling in primary myotubes from WT and PolgA muscle. (**D**,**E**) Cell proliferation was analyzed by EdU labelling and (**F**) cell count. (**G**) Brightfield of 2-d differentiated myotubes. (**H**) Representative blots of dose-response leucine experiment in fully differentiated WT and PolgA myoblasts. (**I**) Quantification of pS6K1 from three independent experiments. (Data represent mean±SEM, *p<0.05 WT (blue line/bars) vs PolgA (red line/bars), #p<0.05 over time determined by two-way ANOVA)

*In vivo*, leucine sensing towards mTORC1 might be altered by the presence of other amino acids or growth factors. To rule out such interference, we cultured myogenic progenitor (MPs) cells from WT and PolgA hind limb muscle to verify proliferation capacity and leucine sensing without the availability of other amino acids. Strikingly, both 5-ethynyl-2’-deoxyuridine (Edu) incorporation and proliferation (*Figure* 5D-F) were reduced in PolgA derived MPs. Furthermore, after differentiation, which was unaffected by PolgA mutation (*Figure* 5G), we subjected myotubes to varying doses of leucine after 60 min of starvation. In contrast to our *in vivo* data, PolgA myotubes showed a strong anabolic resistance to leucine, as indicated by lower pS6K1 at 0.8mM and 5mM leucine, respectively (*Figure* 5H,I). These data show that PolgA muscle progenitors have reduced cell proliferation capacity *in vitro*, potentially due to lower sensitivity to leucine.

### Reduced bone quantity and quality in PolgA mice

As the results of the cross-sectional experiments described above suggest that PolgA mice display clear signs of co-existing osteopenia and sarcopenia from 40 weeks onwards, we aimed to longitudinally track individual mice during the transition from young to frail status in order to gain a better understanding of when exactly and to what extent bone loss (osteopenia) occurs in individual mice. Therefore, we used an *in vivo* micro-CT approach to monitor the dynamic changes in the bone microarchitecture of the 6^th^ caudal vertebrae (CV6) in individual mice between the ages of 20 and 40 weeks (*Figure* 6). The comparison between genotypes showed that at 20 weeks, PolgA and WT had similar trabecular morphometric parameters. Initially, bone volume fraction (BV/TV) and trabecular thickness (Tb.Th) increased in both genotypes, but then started to diverge such that BV/TV and Tb.Th were higher in WT mice from 30 weeks onwards (p<0.05, *Figure* 6A,B). Furthermore, from 30 weeks onwards, Tb.Th continuously increased in WT (+5%, p<0.0001), whereas it did not change in PolgA (-1.3%, p>0.5, *Figure* 6B). No significant differences between genotypes were detected for the trabecular number (Tb.N) and separation (Tb.Sp, *Figure* 6C,D). Nevertheless, Tb.N in WT decreased between 30 and 40 weeks (-2.5%, p<0.05), whereas Tb.N did not change over time in PolgA mice (-0.3%, p>0.05, *Figure* 6C). The reduction in Tb.N together with the increase in Tb.Sp in WT explain the slight decrease in BV/TV in WT mice from 36 weeks onwards. A significant interaction between age and genotype, indicating that the time course developed differently between genotypes, was found for Tb.Th, Tb.N and Tb.Sp.

**Fig. 6.**
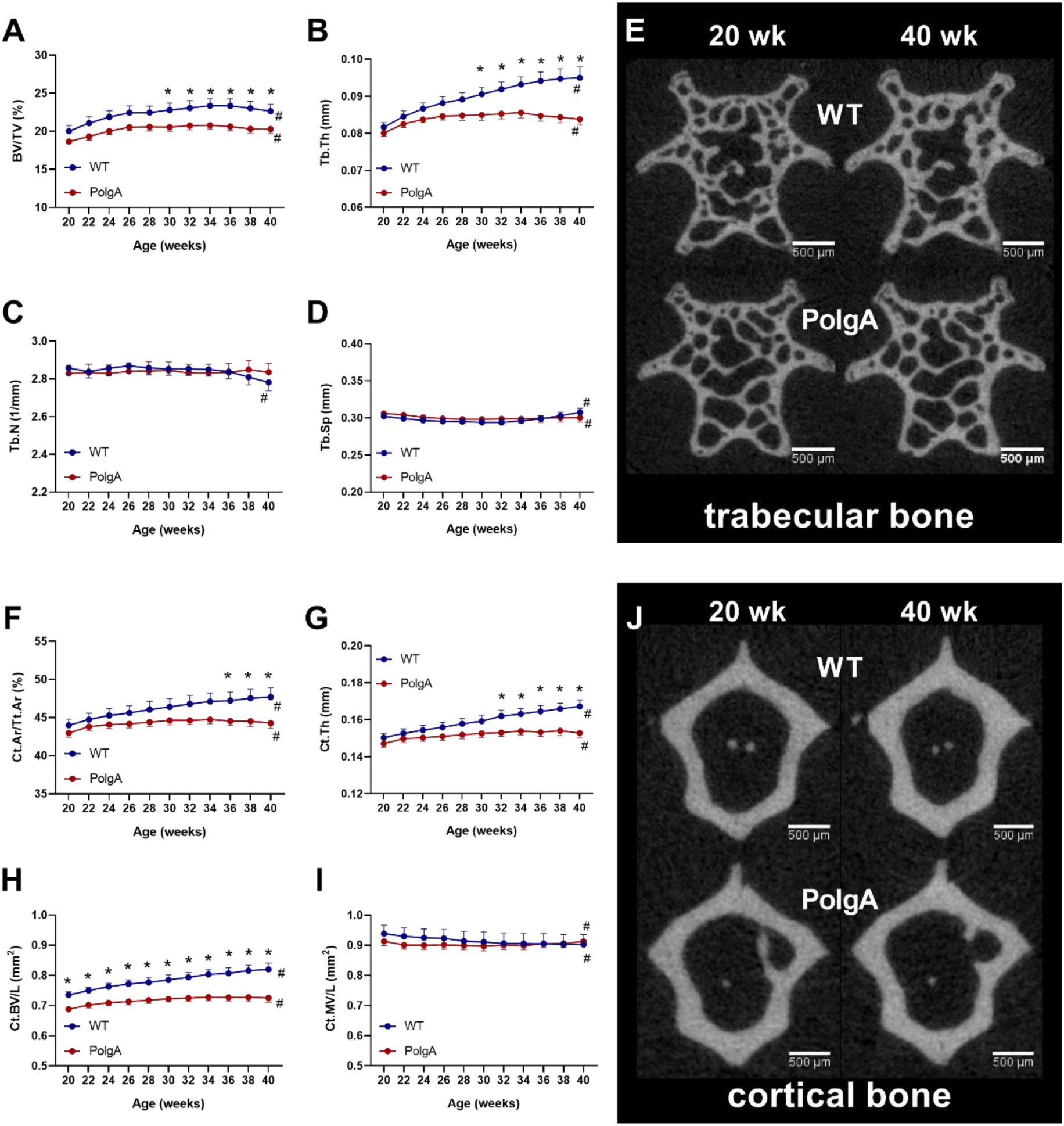
Longitudinal monitoring of trabecular and cortical bone morphometric parameters over 20 weeks: (**A**) bone volume fraction (BV/TV), (**B**) trabecular thickness (Tb.Th), (**C**) trabecular number (Tb.N), (**D**) trabecular separation (Tb.Sp), (**F**) cortical area fraction (Ct.Ar/Tt.Ar), (**G**) cortical thickness (Ct.Th), (**H**) cortical bone volume (Ct.BV) and (**I**) cortical marrow volume (Ct.MV). (**E**,**J**) Bone microstructure (cross-sections) of representative (median) WT (top row) and PolgA (bottom row) mice at 20 (left) and 40 (right) weeks of age. In WT mice, thickening of trabeculae and cortex can be observed, while in PolgA mice, little change can be seen between time points. (Data represent mean±SEM; n=9 WT and n=12 PolgA, *p<0.05 WT (blue line) vs PolgA (red line); #p<0.05 over time determined by linear mixed model and Tukey’s post hoc)

Similar results were obtained when the cortical bone morphometric parameters were analyzed. At 20 weeks, there were no differences between genotypes in cortical area fraction (Ct.Ar/Tt.Ar), Ct.Th and Ct.MV (*Figure* 6F, G, I). Ct.BV was already lower (-6%, p<0.05) in PolgA compared to WT at 20 weeks (*Figure* 6H) with this difference becoming more accentuated up to 40 weeks (-11%, p<0.001). Ct.Th increased over time, but then diverged such that Ct.Th was higher than that of PolgA mice from 32 weeks onwards (p<0.05). This resulted in a higher Ct.Ar/Tt.Ar in PolgA mice from 36 weeks onwards (p<0.05). No differences in Ct.MV were detected between genotypes. A significant interaction between age and genotype was found for Ct.Ar/Tt.Ar, Ct.Th, Ct.BV and Ct.MV. The thickening of the trabecular and cortical structure was also visually apparent from the micro-CT images and can be appreciated when the same cross-section is observed at 20 and 40 weeks (*Figure* 6E,J).

### Reduced bone remodeling in PolgA mice

In addition to monitoring changes in bone morphometry over time, we used *in vivo* micro-CT to quantify dynamic bone formation and resorption parameters, respectively in PolgA and WT caudal vertebrae (*Figure* 7). On average, PolgA mice had lower bone formation rate (BFR, - 23.2%, p<0.05) and bone resorption rate (BRR, -28.8%, p<0.05) compared to WT (*Figure* 7A,B). BFR and BRR changed over time in both genotypes (p<0.001), however the time course did not develop differently between genotypes (interaction effect, p>0.05). Hence, the net remodeling rate (BFR-BRR) was not different between genotypes. The mineral apposition and resorption rate (MAR and MRR, respectively), which represent the thickness of formation and resorption packages were lower (-8.9%, p<0.05 and -20.1%, p<0.001) in PolgA compared to WT (*Figure* 7C,D). MAR did not change over time and showed no significant interaction between age and genotype (*Figure* 6C). On the other hand, there was a significant interaction effect of age and genotype on MRR (p<0.05). Over time, MRR increased in WT, while MRR remained constant in PolgA (*Figure* 7D). The mineralized surface (MS), which represents the surface of formation sites, was lower in PolgA (-11.8%, p<0.0001), whereas the eroded surface (ES) was similar between genotypes (p>0.05) (*Figure* 7E,F). MS and ES changed over time in both genotypes, however the time course did not develop differently between genotypes (interaction effect, p>0.05). In line with lower BFR and BRR observed by micro-CT, bone turnover markers for formation (N-terminal propeptide of type I procollagen (PINP) and resorption (C-terminal telopeptides of type I collagen, CTX-I) were lower in serum from PolgA (-56% and -49%, p<0.05) compared to WT at 40 weeks of age (*Figure* 7H,I). Overall, these results suggest that PolgA mice have reduced bone turnover compared to WT.

**Fig. 7.**
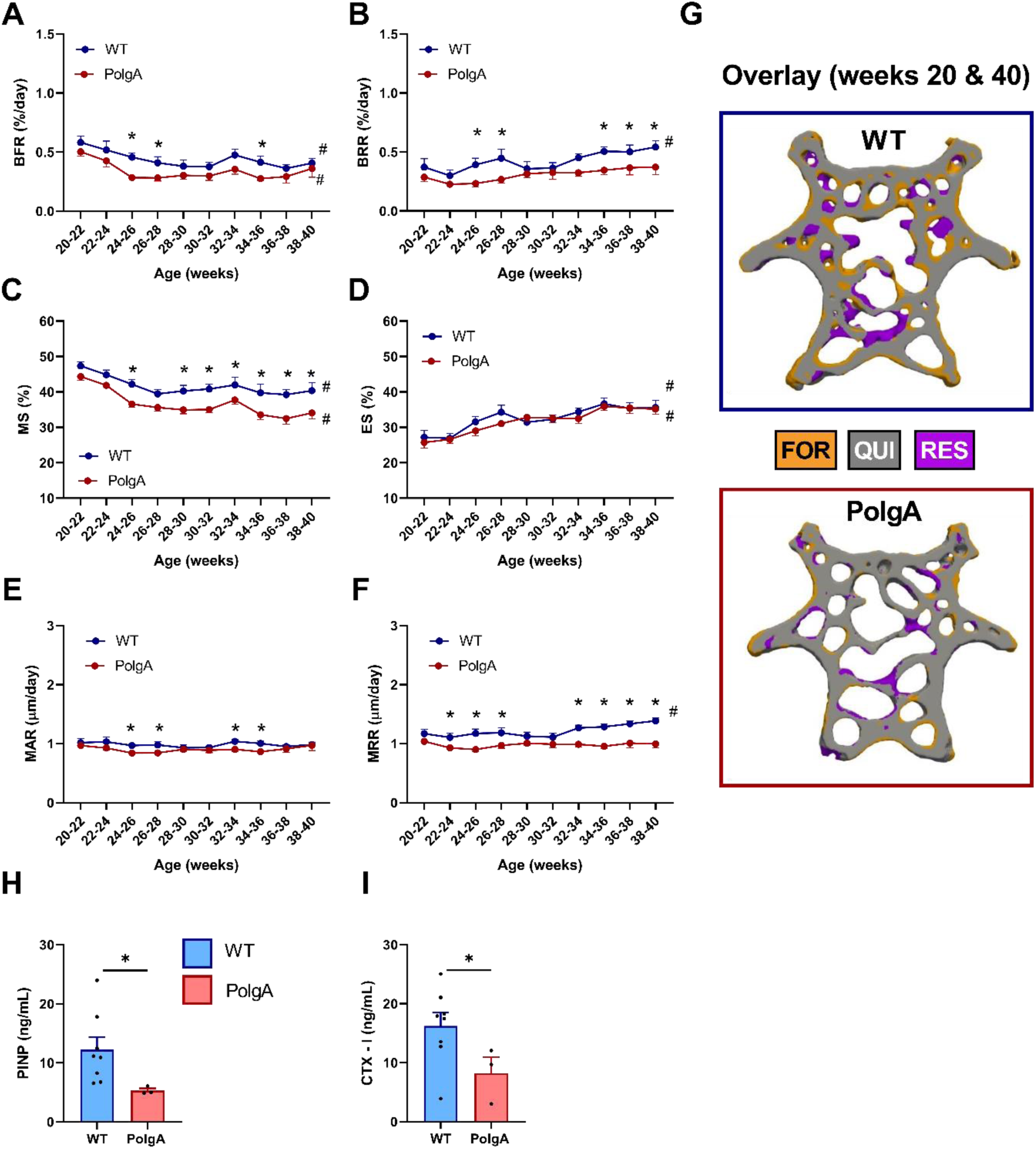
Longitudinal monitoring of dynamic bone morphometry parameters over 20 weeks: (**A**) Bone Formation Rate (BFR), (**B**) Bone Resorption Rate (BRR), (**C**) Mineral Apposition Rate (MAR), (**D**) Mineral Resorption Rate (MRR), (**E**) Mineralizing Surface (MS), and (**F**) Eroding Surface (ES). (Data represent mean±SEM; n=9 WT and n=12 PolgA; *p<0.05 WT (blue line) vs PolgA (red line); #p<0.05 over time determined by linear mixed model and Tukey’s post hoc). (**G**) Overlay of thresholded micro-CT images from 20 and 40 weeks showing sites of formation (orange), quiescence (grey) and resorption (blue) in representative WT and PolgA mouse. Serum bone turnover markers for (**H**) formation (PINP) and (**I**) resorption (CTX-1), respectively at 40 weeks of age. (Data represent mean±SEM; n=8 WT and n=3 PolgA; *p<0.05 WT (blue bar) vs PolgA (red bar) determined by Mann Whitney test)

To assess whether the osteopenic phenotype in PolgA mice could be linked to alterations in mTORC1 signaling, we assessed basal mTORC1 and MAPK signaling in caudal vertebrae at 40 weeks of age. Overall, the expression of downstream mTORC1 and MAPK was lower in vertebrae compared to muscle tissue. Therefore, the bone immunoblots were overexposed in order to observe a signal. Compared to WT, PolgA mice tended to have lower pS6K1 at Thr389 (-69%, p≤0.1) while no significant difference was observed for the direct downstream kinase thereof (RPS6) at Ser235/236 (p>0.05) (*Figure* 8A,B). Basal pSAPK/JNK at Thr183/Tyr185 was lower (-60%, p<0.001) in PolgA compared to WT, while no difference was observed in p44/42 at Thr202/Tyr204 (p>0.05) (*Figure* 8C,D).

**Fig. 8.**
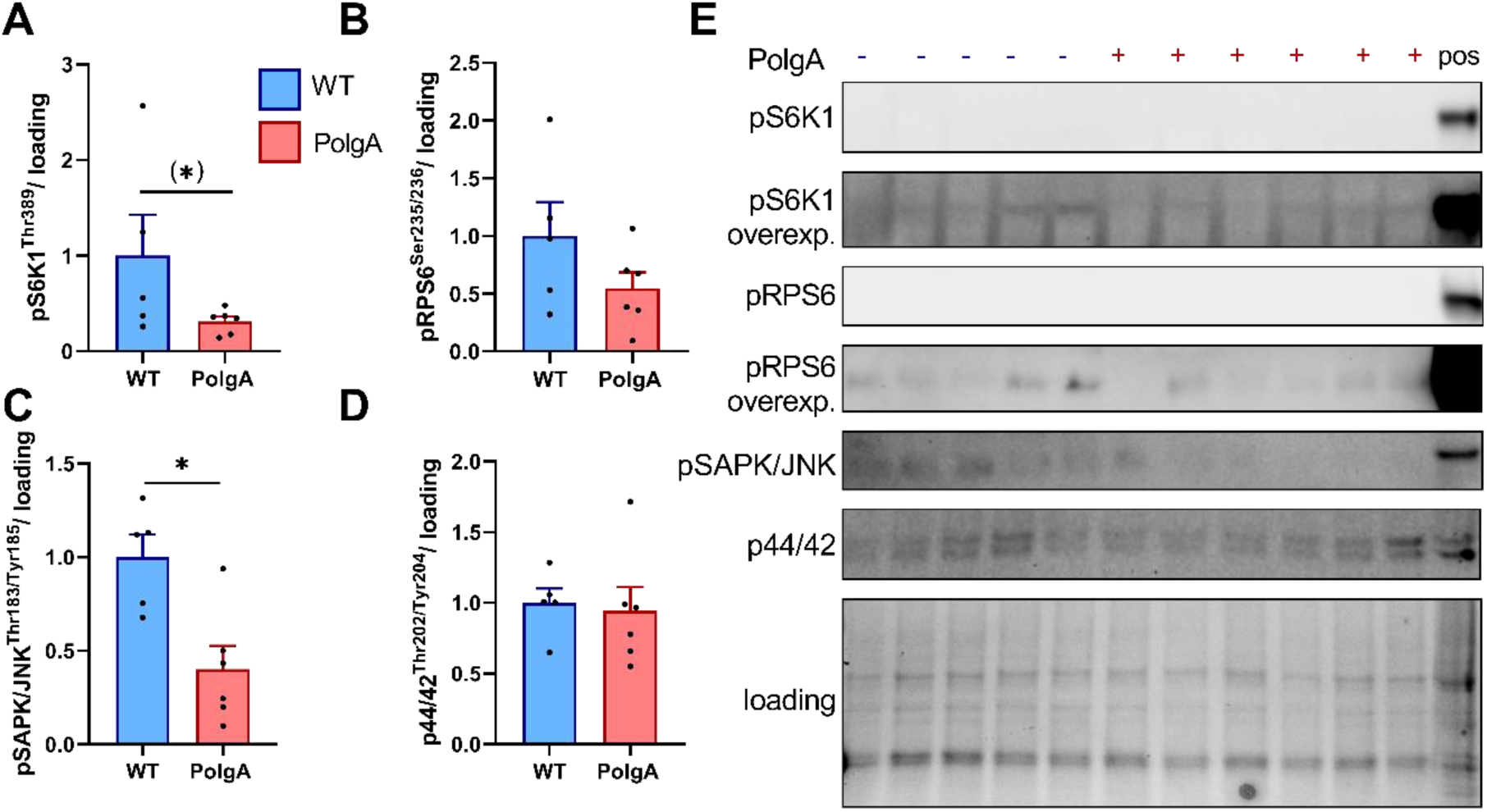
Basal mTORC1 and MAPK signaling in WT and PolgA caudal vertebrae at 40 weeks. PolgA mice showed a tendency towards reduced downstream mTORC1 targets pS6K1 (**A**) and pRPS6 (**B**). Downstream MAPK target pSAPK/JNK (**C**), but not p44/42 MAPK (**D**) was lower in PolgA compared to WT. (**E**) Representative blots with positive control (muscle tissue). Overall signal was low in bone lysates, so pS6K1 and pRPS6 are overexposed. (Data represent mean±SEM, n=5-6/genotype, (*)p≤0.10, *p<0.05 PolgA (red bars) vs WT (blue) by unpaired students t-test)

### Reduced mechanosensitivity of PolgA caudal vertebrae with age

To investigate whether prematurely aged PolgA bones are mechanosensitive, the 6^th^ caudal vertebrae were cyclically loaded 3x/week over 4 weeks using a previously developed loading device [50]. *Figure* 9 shows the absolute values of trabecular bone morphometric parameters at the end of the mechanical loading intervention in young (A,B) and aged (C,D) mice. At young age, loading elicited an anabolic response both in WT and PolgA mice such that BV/TV and Tb.Th were higher (p<0.0001) in the loaded compared to the sham-loaded controls (*Figure* 9A,B). Strikingly, the anabolic effect was maintained with age in WT (p<0.05), whereas PolgA mice did not respond to the mechanical loading intervention, with no differences detected between loaded and non-loaded controls (*Figure* 9C,D).

**Fig. 9.**
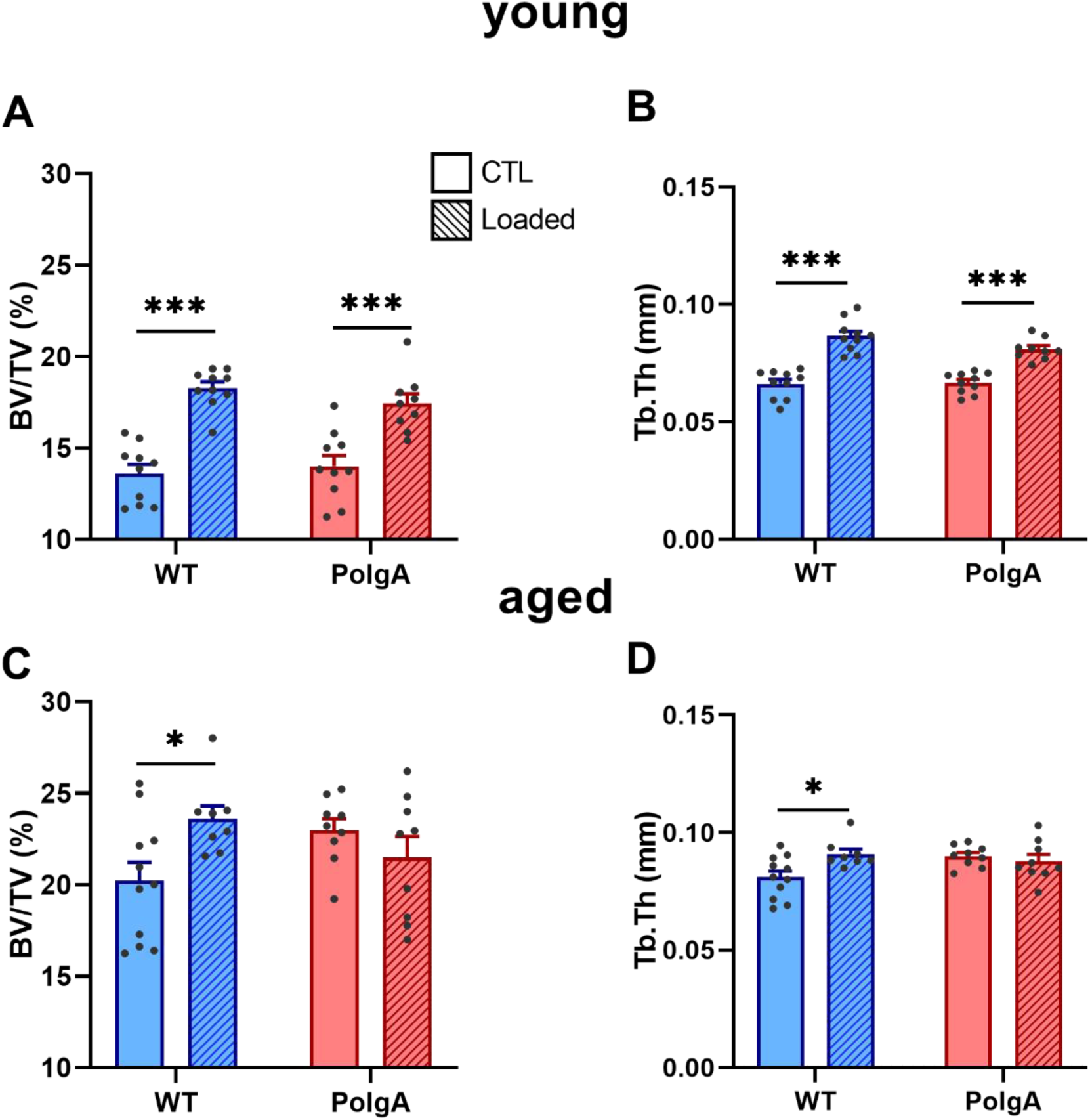
Effect of cyclic mechanical loading on trabecular bone morphometric parameters in PolgA and WT mice at young (top) and old age (bottom): (**A,C**) bone volume fraction (BV/TV) and (**B,D**) trabecular thickness (Tb.Th) at the end of the loading intervention. (Data represent mean±SEM, n=8-11/group; *p<0.05, ***p<0.0001 CTL (circles/solid bars) vs loading (squares/striped bars))

## Discussion

Due to the accumulation of mitochondrial DNA mutations with age, PolgA mice are known to develop multiple signs of aging (alopecia, greying, hearing loss, kyphosis, reduced bone and muscle mass) early in life [34, 35]. Here, we used the clinical mouse FI [29], which was recently reverse translated from the FI in humans [28], to quantify the accumulation of these health deficits over time in individual mice. Furthermore, as the weakening of the musculoskeletal system is a major component of age-related frailty, we used a combination of *in vivo, ex vivo* and *in vitro* techniques to comprehensively characterize the musculoskeletal phenotype of this model. Lastly, the effects of various anabolic interventions on the musculoskeletal system of PolgA mice were assessed.

This study demonstrated that in comparison to their chronologically aged WT littermates, PolgA mice developed many hallmarks of aging over time which collectively resulted in PolgA mice becoming frailer with age. Of the 31 items included in the FI scoring, changes in the integument category were the most striking. Specifically, PolgA mice showed a loss in fur color (from black to grey/brown) and a decline in the coat condition (ruffled fur, ungroomed appearance). Furthermore, they showed signs of enlarged abdomen and kyphosis. At 40 weeks of age (280 days), PolgA mice had mean frailty scores of 0.15±0.02 (mean±SD), whereas WT mice had 0.06±0.04, respectively. A previous study using male C57BL/6J mice reported mean FI values of 0.05±0.05 in young (30-299 days), 0.21±0.04 in middle-aged (300-599 days) and 0.29±0.07 in older mice (above 600 days) [30]. Comparing these values to the ones observed in our study, the (female) WT mice (at 40 weeks < 299 days) fall in the category of young mice, whereas the PolgA mice fall into the category of “middle-aged” mice. Although some studies have reported higher FI scores in female compared to male mice [29, 54], others showed no differences in frailty between sexes [55, 56]; furthermore, it seems that the rate of deficit accumulation is similar between sexes [54]. Nevertheless, future studies in which PolgA mice are aged for a longer period of time (>300 days) would be necessary to investigate the degree of frailty at later stages in life. Our ethical permit for breeding was constrained to a maximum of 46 weeks as previous studies reported median lifespans of PolgA mice ranging from ca. 59 weeks (416 days) [34], 48 weeks (336 days) [35], down to only 36 weeks (250 days) [37]. The discrepancies between studies regarding the lifespan is also apparent for the age at which symptoms of aging first appeared (ranging from 6- [37] to 9 months [34, 35]). As not only homozygous (PolgA^D257A/D257A^), but also heterozygous (PolgA^D257A/+^) mice progressively accumulate mitochondrial DNA point mutations with age, we suspect that the differences between studies resulted from differences in breeding schemes. Since all the mice used in our study were bred according to a standardized breeding scheme, thereby displaying only a single generation of mutation burden, the onset of the accelerated aging phenotype may have been delayed in the present work as compared to other studies. Nevertheless, we observed that PolgA mice became frailer faster than WT, as shown by plotting the natural logarithm of the FI as a function of age and fitting the resulting relationship with a linear function. To our knowledge, we are the first to report the frailty phenotyping in the PolgA mouse model.

In line with increased FI with age, the musculoskeletal phenotype clearly diverged over time between genotypes, with PolgA mice developing multiple signs of osteosarcopenia with age. In agreement with previous reports showing lower muscle weights [34, 36, 37] and reduced femoral bone mineral density assessed by X-ray densitometry [34, 35], phenotyping of the femora and hind limb muscles revealed lower bone and muscle mass as well as cross-sectional area in PolgA mice compared to their WT littermates at 40 and 46 weeks of age. These reductions were not observed at 34 weeks, thus indicating a progressive weakening of the musculoskeletal system in the PolgA mice. As the clinical diagnosis of sarcopenia is not only characterized by reduced muscle mass but also by reduced muscle function and quality, we assessed the forelimb grip-strength, muscle force and intramuscular signaling upon anabolic stimuli. In agreement with clinically diagnosed sarcopenia, PolgA displayed lowered muscle strength, indicated by reduced grip-strength and lower concentric contraction force at different intensities.

One mechanism commonly thought of as a contributor to sarcopenia is the age-related lowered sensitivity to anabolic stimuli. This phenomenon, known as anabolic resistance, has been associated with the reduced acute activation of mTORC1 [13–15], the central protein complex that integrates external stimuli to regulate cell growth [57–60]. Note that the interpretation of the data described in previous studies, which showed increased anabolic resistance upon heavy-load contractions, might be confounded by the fact that the same relative load (% 1 repetition max) was used in old vs. young humans [13]. The lower absolute load in the aged subjects might have caused lower mechanical stress on the muscle, as demonstrated by lower MAPK signaling [61], and thus a dampened increase in mTORC1 signaling. By using the protocol described by O’Neil et al. [44], we could overcome this confounding factor by controlling the load applied to the muscles. Interestingly, although the average force produced during the 60 contractions was similar between WT and PolgA, both MAPK and downstream mTORC1 signaling was remarkably elevated in PolgA mutated muscle. Notably, leucine also activated mTORC1 to a greater extent in 46-week-old PolgA muscle compared to their chronologically aged WT littermates. This is intriguing as eccentric contractions and leucine activate mTORC1 signaling via independent mechanisms [62]. Potentially, increased contraction-induced mechanical signaling is responsible for increased mTORC1 activity in response to contractions, but further research is needed to fully elucidate the mechanisms behind mTORC1 hyperactivity in PolgA muscle. Next to stimulus-induced mTORC1 activity, we also found hyperactive basal mTORC1 in PolgA at 46 weeks of age. In fact, basal mTORC1 hyperactivity has previously also been shown in other models of progeria [63], aging [64] and progeria-like syndromes (i.e., laminopathies) [65, 66]. Furthermore, chronically increased mTORC1 activity has been shown to be sufficient to cause progressive muscle atrophy, fiber damage, fiber death, and muscle weakness in humans and in mice [67]. Therefore, treatment with mTORC1 inhibitors has been proposed as a potential therapeutic approach to rescue elevated mTOR signaling in these models [65–67].

*In vivo*, the systemic availability of amino acids/nutrients and differences in the local niche might have interfered with hyperactive mTORC1 signaling in PolgA. To rule this out, we used an *in vitro* approach to study the activation of mTORC1 in MPs that were stimulated with identical concentrations of leucine, without the availability of other amino acids. We were able to demonstrate that PolgA primary MPs had substantial defects in proliferative capacity and were completely insensitive to leucine with respect to mTORC1 anabolic signaling. Thus, it seems that mitochondrial dysfunction per se is sufficient to induce impairments in muscle stem cell mTORC1 signaling, regardless of exposure to the local niche. The differences in downstream mTORC1 signaling upon anabolic signals between whole muscle and their respective stem cells are puzzling, but can potentially be induced by the higher accumulation of mitochondrial damage under proliferative *in vitro* conditions when compared to post mitotic differentiated muscle tissue that develops apoptosis at a later stage [34]. Nonetheless, these data warrant caution in the translation of MP-derived experimental data to the *in vivo* muscle setting and suggest that the accumulation of mitochondrial damage is tissue specific and can profoundly change the mTORC1 response to leucine.

With respect to the evaluation of the bone phenotype, this study is to the best of our knowledge, the first to use standardized micro-CT approaches to comprehensively characterize the bone phenotype of the PolgA mouse model [38, 39, 48]. Albeit still considered the clinical gold standard for assessing bone mineral density in humans, X-ray densitometry, which was previously used to evaluate the PolgA bone phenotype [34, 35], is confounded by the size and positioning of the bone, and furthermore, does not provide any information on the bone microarchitecture [68, 69]. Nevertheless, the results of this study did recapitulate the clear reductions measured in femoral bone mineral density in PolgA mice with age [34, 35]. Note that the accuracy of the aforementioned phenotyping of the femora, both in the previous as well as in the current study, is limited by the cross-sectional study design. This type of study design cannot account for the intrinsic initial variability between animals at baseline [70, 71], thereby making it impossible to determine the amount of bone lost in individual mice. We therefore used an established *in vivo* micro-CT approach to monitor the spatio-temporal changes in bone micro-architecture in the caudal vertebrae of individual mice during the transition from healthy to frail state. Multiple bone morphometric parameters diverged over time, from being similar at 20 weeks to lower in PolgA mice with age. Interestingly though, this difference did not arise due to significant bone loss in PolgA mice, but rather in the failure to achieve normal peak bone mass. We and others have previously reported the absence of age-related bone loss in caudal vertebrae [72, 73]. Hence, mouse caudal vertebrae may not be optimal for investigating age-related bone loss. As the bone morphometric parameters in the femora declined with age in PolgA mice, this study suggests that age-related changes in bone micro-architecture differ depending on the skeletal site that is analyzed. Indeed, a more severe age-related deterioration of bone microarchitecture in long bones compared to lumbar vertebrae has previously been reported [71, 72, 74]. In a cross-sectional study performed by Glatt et al., the cross-sectional area of the lumbar and caudal vertebrae of female C57BL/6 mice increased by 25% between the age of 2 and 20 months, while the femoral cross-sectional area declined by 3% [71]. Furthermore, using *in vivo* micro-CT, we and others have previously shown differences in age-related changes in the lumbar and caudal vertebrae as compared to the tibiae of C57BL/6 mice [73, 74]. Both studies showed continuous declines in cortical bone of the tibiae, while no changes or even increases were observed in the lumbar and caudal vertebrae, respectively [73, 74]. We therefore assume that the osteopenic phenotype in PolgA mice is stronger in long bones compared to vertebrae. Hence, monitoring the long bones of PolgA mice by *in vivo* micro-CT may be beneficial in future studies. Beyond enhanced phenotyping, the current study capitalized on the ability of *in vivo* micro-CT to provide information on the dynamic coupling between bone resorption and formation. Compared to WT littermates, PolgA had lower bone remodeling activities as shown by reduced bone formation and resorption rates, with no differences in the net remodeling rate. These results were confirmed by lower bone turnover markers measured in PolgA serum compared to WT. Similarly, senile osteoporosis in humans is characterized by low bone turnover, as opposed to the high bone turnover rates (higher resorption activities) observed during postmenopausal osteoporosis. Despite the advantages of *in vivo* micro-CT, the anesthesia, cumulative radiation, and stress associated with repeated CT measurements could have an effect on the bone morphometry and the well-being of the animals [75–79]. However, as effects of radiation are of greater concern in young, growing mice, and furthermore, have been shown to be independent of surgical treatments (e.g., ovariectomy) [75], we suspect that any effects of radiation would be similar in WT and PolgA mice. Nevertheless, we compared the bone remodeling activities of PolgA and WT mice used in this study (subjected to 11 *in vivo* measurements) with those of corresponding age-matched controls that received only 2 *in vivo* scans at 38 and 40 weeks, respectively, and saw no statistical differences between the groups. Furthermore, we did not observe any negative effects of any of these variables on the well-being of the animals.

Finally, after having established that PolgA mutation altered the acute sensitivity to anabolic stimuli in muscles and their progenitors, we investigated the responsiveness of PolgA bones to a previously established 4-week mechanical loading regime [49]. Interestingly, prematurely aged PolgA mice were not responsive to cyclic loading of the caudal vertebrae over 4 weeks, whereas loading had beneficial effects in WT mice. Although there are controversial reports on the maintenance of bone mechanosensitivity with age in mice [23–26], these results conflicted with our previous observation that caudal vertebrae remain mechanosensitive to cyclic mechanical loading with age [27]. Interestingly, in that study, 82-week-old C57BL/6J mice showed a greater anabolic response compared to the 52-week-old mice; hence, it is possible that even older PolgA mice would respond differently than the ones used in this study. Furthermore, the load in this study was not adjusted to account for differences in the initial bone volume fraction between PolgA and WT mice. An individualized loading approach, which ensures that all bones are subjected to the same mechanical strain, could yield differences in the mechanosensitivity of the PolgA and WT mice. Nevertheless, adapting the loading for the lower bone mass in the PolgA would decrease the stimulus and therefore, we do not expect a positive response at this even lower stimulus. Nonetheless, further research is therefore needed to elucidate the mechanosensitivity of PolgA bones. Of further notice was that BV/TV and Tb.Th declined over time in the aged sham-loaded WT mice, whereas loading was able to reduce this bone loss. As our *in vivo* micro-CT monitoring of the caudal vertebrae did not show any bone loss up to 40 weeks of age (*Figures* 5 and 6), we suspect that the bone loss occurring in the loaded WT mice was due to the surgical insertion of the pins required for loading.

Interestingly, this decline in trabecular bone was not observed in the PolgA mice, thus explaining the high BV/TV and Tb.Th in the sham-loaded PolgA mice at end-point. We have previously observed similar bone loss in 52-week-old mice with loading being able to reduce this bone loss [27]. Similar to our current study though, the 15- and 82-week-old mice did not show any bone loss. It is possible that the prematurely aged PolgA mice behave similarly to the 82-week-old mice of that study. Nevertheless, the fact that PolgA bones lose mechano-responsiveness to loading with age, while PolgA muscles show increased acute response to mechanical stimuli is intriguing. Along the same lines were the contrasting results in terms of basal mTORC1 and MAPK signaling, where basal downstream mTORC1 and MAPK signaling was almost absent in PolgA mutated bone, while it was hyperactive in PolgA muscle. Tissue-specific differences in mTORC1 signaling have been shown previously [80]. Genetic reduction of mTOR expression had beneficial effects on neurological and muscle function (e.g., rotarod, grip-strength) in aging mice, whereas declines in bone volume and immune function were accelerated [80]. Moreover, the pharmacological inhibition of mTORC1 signaling has been shown to impair osteogenic [81, 82] and osteoclast [83] differentiation *in vitro* as well as induce trabecular bone loss *in vivo* [84], while rapamycin treatment reduced muscle fiber loss in aging mice [67]. Hence, although treatments with mTOR inhibitors could be beneficial for long-term muscle health, this might not be the case for bone health. Future studies are required to better understand discrepancies and potential interactions in mTORC1 signaling between muscle and bone, and whether such differences can be linked to altered mechanosensitivity with age.

There are a number of limitations to our study that should be mentioned. Firstly, the acute response to anabolic stimuli in the muscle tissue cannot be directly compared to the long-term loading regime (over 4 weeks) to which the bone tissue was subjected. Physiological resistance training protocols leading to skeletal muscle hypertrophy in mice are difficult to achieve. The protocols, which are available such as voluntary resistance running [85] and high intensity interval running [86] might be cumbersome due to a reduced exercise capacity in PolgA [87]. Nevertheless, both voluntary [19] and forced [37] long-term endurance exercise regimes have been shown to improve the progeroid phenotype and locomotor behavior in PolgA mice. Hence, future studies should investigate the potential of longer-term resistance exercise training in alleviation of mTORC1 hyperactivity in (mouse models of) progeria. To be noted though is that the above mentioned exercise regimes were applied already at young (10-12 weeks) up to old age thereby making it unknown whether they would have the same effect in older PolgA mice already displaying signs of aging. In this respect, future studies subjecting PolgA caudal vertebrae to cyclic loading regimes throuoghout life (i.e, from young until old age) would be interesting to assess whether the reduction in bone mechanosensitivity with age could be prevented.

A further limitation of our study was that the evaluation of the bone remodeling activities was mainly based on time-lapsed micro-CT analysis, whereas two-dimensional (2D) histomorphometry using fluorescent labels would allow for the assessment of bone formation and mineralization at an even higher resolution [88]. However, owing to the lack of an appropriate marker, bone resorption can only be estimated from the number of osteoclasts. In this regard, the non-destructive nature of time-lapsed micro-CT imaging provides a major advantage as both bone formation as well as bone resorption can be assessed longitudinally in individual mice [25, 39], thereby reducing the number of animals needed for experiments. Nevertheless, we and others have previously shown high correlations between dynamic histomorphometric parameters quantified by micro-CT and conventional histomorphometry, respectively [25, 39, 89]. A further advantage of micro-CT, however, is that bone histomorphometric parameters can be assessed in the entire 3D compartment rather than in a limited number of 2D histological sections. This not only reduces labor-intensive processing of the samples but also reduces inter-and intra-observer errors associated with the analysis thereof [90, 91]. The final determining advantage for using *in vivo* micro-CT in this particular study though was the long-term period of 20 weeks over which bone remodeling activities were assessed. Due to the resorption of fluorochrome labels, 2D histomorphometry requires very short intervals (2-3 days) between marker injections [92], and hence, is not well suited for long-term studies.

In conclusion, we show that PolgA mice develop multiple hallmarks of aging, such as reduced bone remodeling and muscle mass early in life, which collectively can be quantified using the mouse FI. We further demonstrate acute mTORC1 hyperactivity in PolgA muscle upon anabolic signals, which is related to diminished satellite cell proliferation. By mimicking many aspects of osteosarcopenia, the PolgA mouse provides a powerful model that facilitates our understanding of the relationship between muscles and bones, and also between the aging musculoskeletal system and frailty.

## Acknowledgements

We gratefully acknowledge Dr. Ilaria Bellantuono for her guidance and inputs for the quantification of the clinical mouse frailty index. We acknowledge Susanne Friedrich for performing the force-grip assessments. The authors certify that they comply with the ethical guidelines for publishing in the Journal of Cachexia, Sarcopenia and Muscle: update 2017 [96].

## Funding

This manuscript is based upon work supported by the Swiss National Science Foundation (SNF IZCNZ0-174586), the European Cooperation in Science and Technology (COST Action BM1402: MouseAGE) and the European Research Counsil (ERC Advanced MechAGE ERC-2016-ADG-741883).

## Author contributions

ACS and GDH designed the experiments, collected the data and wrote the manuscript, GAK designed the experiments, collected the data and reviewed the manuscript, ME collected the data and reviewed the manuscript, ESW provided support establishing methods and reviewed the manuscript, RM and KDB designed the experiments and wrote the manuscript.

## Competing Interests

All authors declare that they have no conflicts of interest.

## Notes

#### Summary of Updates

Small modifications in manuscript main text and 2 figures. Supplemental files updated.

