## Supplementary Material and Methods for "Hallmarks of frailty and osteosarcopenia in prematurely aged PolgA^D257A/D257A^ mice"

**The PDF file includes:**

Supplementary Materials and Methods

Table S1: Animal experiments: Overview and analysis details.

Table S2: Reagents and antibodies used for immunoblotting.

Table S3: Cortical bone morphometry of femora.

**Supplementary Materials and Methods**

**Breeding scheme**

As both heterozygous (PolgA$^{D257A/+}$) and homozygous (PolgA$^{D257A/D257A}$) mice have progressive accumulation of mitochondrial DNA point mutations, specific breeding considerations were taken into account while expanding the colony. Specifically, as paternal mitochondrial DNA is actively eliminated following fertilization in mice, the (undesirable) accumulation of mutations in the germline was minimized by mating heterozygous male mice (PolgA$^{D257A/+}$) with C57Bl/6J inbred females (Charles River Laboratories, Sulzfeld, Germany). The thus obtained heterozygous (PolgA$^{D257A/+}$) females and males (originating from a wild type (WT) C57BL/6J mother) were crossed (age 7-9 weeks) to generate homozygous (PolgA$^{D257A/D257A}$, referred to as PolgA) and WT littermates (PolgA$^{+/+}$, referred to as WT) with only a single generation of mutation burden.

**Mouse genotyping**

The presence of the PolgA knock-in mutation was confirmed by extracting DNA from ear clips (Sigma-Aldrich, KAPA Express Extract, KK7103) followed by qPCR (Bio-Rad, SsoAdvanced Universal SYBR Green Supermix, 1725272) and melt curve analysis. The primers used for genotyping (5' to 3'; Rev Common: AGT AGT CCT GCG CCA ACA CAG; Wild type forward: GCT TTG CTT GAT CTC TGC TC; Mutant forward: ACG AAG TTA TTA GGT CCC TCG AC) were recommended by the Jackson Laboratory.

**Complex IV Enzyme Activity**

In order to confirm that the premature aging phenotypes were associated with mitochondrial dysfunction, we measured the activity of complex IV enzyme in *m. gastrocnemius* (GAS) using the Complex IV Rodent Enzyme Activity Microplate Assay Kit (ab109911, Abcam). Briefly, GAS was homogenized using a tissue homogenizer (Omni THg, Omni International) and protein concentration was determined using the DC assay protein method (Bio-Rad). 50 µg of proteins in 200 µL assay solution were loaded into each well of 96 well plates coated with COX specific antibodies, and incubated at room temperature for 3 hours. The oxidative capacity of COX in the presence of cytochrome C was monitored by absorbance at 550 nm at 30°C for 120 minutes using a photospectrometer (Spark, TECAN), with a measurement interval of 1 minute. Rates of oxidation were calculated by using absorbance values during the time when the decrease of OD values was linear. We found that COX IV activity in PolgA muscles was lower compared to WT muscles both at 40 (-19%, $p<0.0001$) and 46 weeks (-24%, $p<0.0001$), thus confirming that the mice used in this study had the same phenotype as those previously reported [36-37, 93].

**Muscle progenitor isolation**

Primary muscle progenitor cells (MPs) were extracted according to an adapted protocol described by Gorski et al. [94]. In brief, muscle tissue was digested with 2mg/mL collagenase type II (Thermo Scientific Scientific) in collagenase buffer 2% Fetal Bovine Serum (FBS) (Invitrogen) in Hanks Balance Salt Solution (HBSS) (Invitrogen) for 1h at 37°C. The cell pellet was filtered using 40 and 100µm cell strainers. A heterogeneous cell population was purified from fibroblasts by serial pre-plating (1.5h) at day 0, 4 and 10 after cell extraction. Muscle progenitors were cultured on matrigel-coated dishes (Corning, 1/25 dilution) in medium containing 40% F10 nutrient mixture (Invitrogen, 40% low-glucose DMEM (Invitrogen), 20% FBS, 1% Penicillin-Streptomycin (P/S) (Invitrogen) and 5 ng/mL basic fibroblast growth factor (Thermo Fischer Scientific).

**Muscle fiber cross-sectional area determination**

For histology and morphometry of skeletal muscle, 10 µm thick cryosections of TA muscles were dried and washed for 5 min in PBS supplemented with 0.05% triton and subsequently incubated with wheat germ agglutinin Alexa Fluor 647 (1:250, Thermo Fischer Scientific) for 1h. Slides were mounted after a 3x 5min wash in PBS, sealed with glass cover slips and imaged with an epifluorescent microscope (Zeiss Axio observer Z.1) at 10x. Fiber cross-sectional area was automatically determined with a Muscle J plugin for Image J software [95].

**Table S1:** Animal experiments: Overview, setup, and analysis details

| Experiment | Experiment details |
|---|---|
| **1. Frailty and musculoskeletal phenotyping in PolgA vs. WT:** | Total sample size: n=88 female mice<br>4 groups with each group:<br>WT: n=10<br>PolgA: n=12 |
| **- Cross-sectional comparisons at 34, 40 and 46 weeks** | 3 groups<br>*in vivo* measurements:<br>- Frailty Index (Fig. 1)<br>- Forelimb Grip-Strength (Fig. 2)<br>*ex vivo* measurements:<br>- Micro-CT of femora (Fig. 1)<br>- Muscle weights & forces (Fig.1 & 2)<br>- Eccentric Contractions (Fig. 3 & 4) |
| **- Longitudinal monitoring of frailty and bone morphometry** | 1 group<br>- Frailty Index (Fig. 1) measured at 34, 38 and 40 weeks<br>- Bi-weekly *in vivo* micro-CT of caudal vertebrae (Fig. 6 & 7)<br>- Western blots from caudal vertebrae (Fig. 8) |
| - Bone turnover markers from serum at 40 weeks | Total sample size: n=11 female mice<br>WT: n= 8<br>PolgA: n=3<br>(Fig. 7) |
| **2. Leucine administration** | Total sample size: n=18 female mice (intervention at 46 weeks)<br><br>WT: n=3 CTL and n=6 LEU<br>PolgA: n=3 CTL and n=6 LEU<br><br>- Leucine administration (Fig. 5) |
| **3. Cyclic mechanical loading of caudal vertebrae** | Total sample size: n=80 female mice<br><br>Young mice (intervention start at 15 weeks):<br>WT: n=10 CTL and n=10 Loaded<br>PolgA: n=10 CTL and n=9 Loaded<br><br>Aged mice (intervention start at 38 weeks):<br>WT: n=11 CTL and n=8 Loaded<br>PolgA: n=9 CTL and n=9 Loaded |

| | |
|---|---|
| | - Cyclic mechanical loading of 6th caudal vertebrae (over 4 weeks)<br>- *In vivo* micro-CT of caudal vertebrae (Fig. 9) |

**Table S2:** Antibodies used for Immunoblotting

| **Reagents** | |
|---|---|
| Lysis buffer | (1:10, w/v) [50mM Tris-HCl pH 7.0, 270mM sucrose, 5mM EGTA, 1mM EDTA, 1mM sodium orthovanadate, 50mM glycerophosphate, 5mM sodium pyrophosphate, 50mM sodium fluoride, 1mM DTT, 0.1% Triton-X 100 and 10% protease inhibitor] |
| TBST | TBS (1:10, w/v) [24.23g Trizma HCl, 80.06g NaCl in 800ml ultra-pure water pH 7.6, topped up to 1 L] with 1ml of Tween |
| **Primary antibodies 1:1000** | |
| p-p70S6K$^{Thr389}$ | 9234, Cell Signaling |
| p-RPS6$^{Ser235/236}$ | 2211, Cell Signaling |
| p-SAPK/JNK$^{Thr183/Tyr185}$ | 9251, Cell Signaling |
| p-p44/42 MAPK (Erk1/2)$^{Thr202/Tyr204}$ | 9101, Cell Signaling |
| p-ULK$^{Ser757}$ | 6888, Cell Signaling |
| p-mTOR$^{Ser2448}$ | 5536, Cell Signaling |
| **Secondary antibody 1:5000** | |
| Anti-Rabbit IgG, HRP-linked Antibody | 7074, Cell Signaling |

**Table S3: Cortical bone morphometry of femora. Two-dimensional cortical bone morphometric parameters in WT and PolgA mice at 34, 40 and 46 weeks, respectively. (Data represent mean±SD, ** p<0.01, *** p<0.0001 WT vs PolgA at given age)**

| | WT | | | PolgA | | |
|---|---|---|---|---|---|---|
| **Parameter** | **34 weeks** | **40 weeks** | **46 weeks** | **34 weeks** | **40 weeks** | **46 weeks** |
| **Ct.Ar ($mm^2$)** | 0.778 ±0.027 | 0.780 ±0.038 | 0.796 ±0.066 | 0.755 ±0.031 | 0.679 ±0.054*** | 0.664 ±0.053*** |
| **Tt.Ar ($mm^2$)** | 1.684 ±0.05 | 1.702 ±0.066 | 1.820 ±0.061 | 1.633 ±0.075 | 1.615 ±0.084** | 1.694 ±0.074** |
| **Ct.Ar/Tt.Ar (%)** | 46.208 ±1.686 | 45.802 ±1.018 | 43.755 ±2.958 | 46.316 ±2.099 | 42.015 ±2.336** | 39.064 ±2.576*** |
| **Ct.Th (mm)** | 0.196 ±0.008 | 0.199 ±0.006 | 0.195 ±0.016 | 0.197 ±0.008 | 0.177 ±0.013*** | 0.166 ±0.012*** |
| **Ps.Pm (mm)** | 4.680 ±0.092 | 4.701 ±0.100 | 4.847 ±0.086 | 4.581 ±0.118 | 4.564 ±0.134** | 4.654 ±0.105*** |
| **Ec.Pm (mm)** | 3.437 ±0.109 | 3.441 ±0.089 | 3.638 ±0.13 | 3.382 ±0.142 | 3.482 ±0.126 | 3.648 ±0.1 |
| **Length (mm)** | 15.996 ±0.227 | 16.29 ±0.122 | 16.269 ±0.206 | 15.922 ±0.283 | 15.916 ±0.298** | 16.085 ±0.222** |
